# Near 100% efficient homology-dependent genome engineering in the human fungal pathogen *Cryptococcus neoformans*

**DOI:** 10.1101/2025.02.14.638277

**Authors:** Matthew J. Nalley, Sanjita Banerjee, Manning Y. Huang, Hiten D. Madhani

## Abstract

We recently described CRISPR/Cas9-based short homology-dependent genome engineering in the human fungal pathogen *Cryptococcus neoformans*, a haploid budding yeast that is the most common cause of fungal meningitis and an emerging model organism. This was achieved by electroporation of strains stably expressing a codon-optimized Cas9 with two separate DNA molecules, one encoding a selectable marker flanked by short homology arms and a second encoding a sgRNA under the control of the U6 snRNA promoter. However, the efficiency of desired homology-dependent repair relative to undesired non-homologous end-joining (NHEJ) events can be low and variable. Here, we describe methods and strains enabling extremely efficient (∼99%) homology-dependent genome editing in *C. neoformans*. This high-efficiency method requires two manipulations. First, we placed the sgRNA-expressing segment into the marker-containing DNA flanked by targeting homology; thus, only a single DNA molecule is introduced into cells. Second, we used a strain mutant for the non-homologous end-joining factor Ku80 (encoded by *YKU80*). We also report the engineering of a *yku80::amdS* mutant strain harboring an insertion mutation that can be removed scarlessly via recombination between direct repeats. This enables the functional restoration of *YKU80* after homology-dependent genome editing via selection against the *amdS* marker using fluoroacetamide. This approach minimizes documented drawbacks of using Ku-defective strains in downstream experiments. Finally, we describe a plasmid series that enables rapid cloning of sgRNA-marker constructs for genomic manipulation of *C. neoformans*, including gene deletion and C-terminal tagging. These methods, strains, and plasmids accelerate the genomic manipulation of *C. neoformans*.

## Introduction

The human fungal pathogen *Cryptococcus neoformans* is a haploid basidiomycete budding yeast with a complete sexual cycle and excellent animal models for infection (CHUN AND MADHANI 2010). *C. neoformans* is clinically significant, being responsible for >110,000 deaths from meningitis in immunocompromised patient populations (RAJASINGHAM *et al*. 2022). Consequently, it sits atop the W.H.O. ranked list of priority fungal pathogens(FISHER AND DENNING 2023).

Methods to transform *Cryptococcus neoformans* have improved significantly since biolistic (“gene gun”) transformation was introduced over three decades ago (TOFFALETTI *et al*. 1993). The implementation of CRISPR-Cas9 has made electroporation a feasible alternative to biolistics (ARRAS *et al*. 2016; WANG *et al*. 2016; WANG *et al*. 2017; FAN AND LIN 2018; WANG 2018; LIN *et al*. 2020). In one approach called TRACE, co-electroporation of three separate linear DNA molecules is utilized: one encoding a selectable marker flanked by targeting homology, a second encoding Cas9 codon-optimized for expression in the nematode worm *C. elegans* and harboring *C. elegans* introns, and a third encoding a sgRNA whose expression is driven by the U6 promoter. This approach still required long homology arms for genome editing (FAN AND LIN 2018; LIN *et al*. 2020). By codon-optimizing Cas9 for *C. neoformans*, including a *C. neoformans* intron in the Cas9-encoding segment, and integrating this optimized Cas9 into the genome, we previously described short homology-directed repair (HDR) (HUANG *et al*. 2022). This approach allows for ∼50 bp homology arms, easily added by PCR, eliminating the need for long 1kb homology arms. Although these advances have greatly facilitated genome manipulation, we sought to improve the efficiency of HDR, which remains low relative to nonhomologous end-joining (NHEJ).

While *C. neoformans* has at least two DNA repair pathways for repairing double-stranded breaks induced by Cas9-sgRNA complexes (NHEJ and HDR), the dominant pathway is NHEJ. Thus, using the state-of-the-art system, many colonies must typically be screened to obtain the desired genetic modification. Previous work has indicated that the NHEJ pathway can be inhibited using the small molecule W7 to increase the proportion of transformants that utilize the HDR pathway (ARRAS *et al*. 2016). However, results were mixed, and W7 was not beneficial in some experiments (ARRAS AND FRASER 2016; HUANG *et al*. 2022). Other studies have shown that essential components of the NHEJ pathway, Ku70 and Ku80 (CRITCHLOW AND JACKSON 1998; VALENCIA *et al*. 2001) (encoded by *YKU70* and *YKU80*), could be mutated to reduce NHEJ (GOINS *et al*. 2006; LIN *et al*. 2015). Although HDR efficiency increased in this context, it did not approach 100%. Conversely, others reported no HDR bias in these mutant backgrounds based on no significant change in the number of transformants obtained in NHEJ-deficient backgrounds. However, the transformants obtained were not analyzed to determine whether they were produced by NHEJ versus homologous recombination (FAN AND LIN 2018). Complicating the use of NHEJ-defective mutants are reports that Ku-defective mutants in *C. neoformans* exhibit increased mutation rates (JUNG *et al*. 2021), and that a *yku80Δ* mutant impacts virulence and stress adaptation (LIU *et al*. 2008).

Here, we describe methods and strains that reliably enable extremely efficient genome editing in *Cryptococcus neoformans*. We show that fusing the sgRNA-encoding DNA to a selectable marker, flanking this construct with short homology arms, and then electroporating a Ku-deficient strain constitutively expressing a codon-optimized Cas9 reliably yields >99% efficient homologous recombination after selection. To address the concerns arising from using NHEJ-deficient strains, we generated a strain, the “*yku80*-blaster” strain, harboring a *reversible* mutation in *YKU80* such that a wild-type *YKU80* allele can be regenerated using a counter-selectable marker that selects for recombination between direct repeats flanking the marker. The *yku80*-blaster strain provides the benefits of near-perfect HDR efficiency and the ability to restore Ku80 function, thereby minimizing the consequences of Ku deficiency. Finally, we describe a series of plasmids enabling facile generation of sgRNA-marker constructs useful for both genomic deletion and C-terminal protein tagging. These methods, plasmids, and the Cas9-expressing, *yku80*-blaster strain accelerate the genetic manipulation of *C. neoformans*.

## Results

We previously reported genome engineering of *Cryptococcus neoformans* via homology-directed repair (HDR) with short homology arms using a strain constitutively expressing a codon-optimized Cas9 (HUANG *et al*. 2022). We used the TRACE methodology (LIN *et al*. 2020) to transform strains with 1) a marker molecule containing homology to the desired recombination sites in the genome and 2) a separate molecule encoding a sgRNA driven by the U6 snRNA promoter. We reported homology-dependent recombination frequencies ranging from 9% to 44% of all transformants (ERPF *et al*. 2019; HUANG *et al*. 2022). Frustratingly, this efficiency can be much lower for some genes, especially large genes, requiring the screening of very large numbers of transformants to identify transformants harboring the desired recombination events (unpublished observations).

### Mutation of *YKU80* enforces HDR

Driven by our need to improve the efficiency of HDR, we revisited the impact of *YKU80* mutations. As mentioned above, the *number* of transformants with a marker gene flanked by homology does not change dramatically in strains lacking NHEJ, but this does not in itself reveal whether transformants obtained after selection for the marker gene are produced by NHEJ or HDR (Fig. 1a). Therefore, we transformed a *yku80*Δ strain with a DNA encoding single guide targeting the *ADE2* gene and a second DNA encoding either 1) a marker lacking homology arms or 2) a marker with short homology arms.

**Figure 1.**
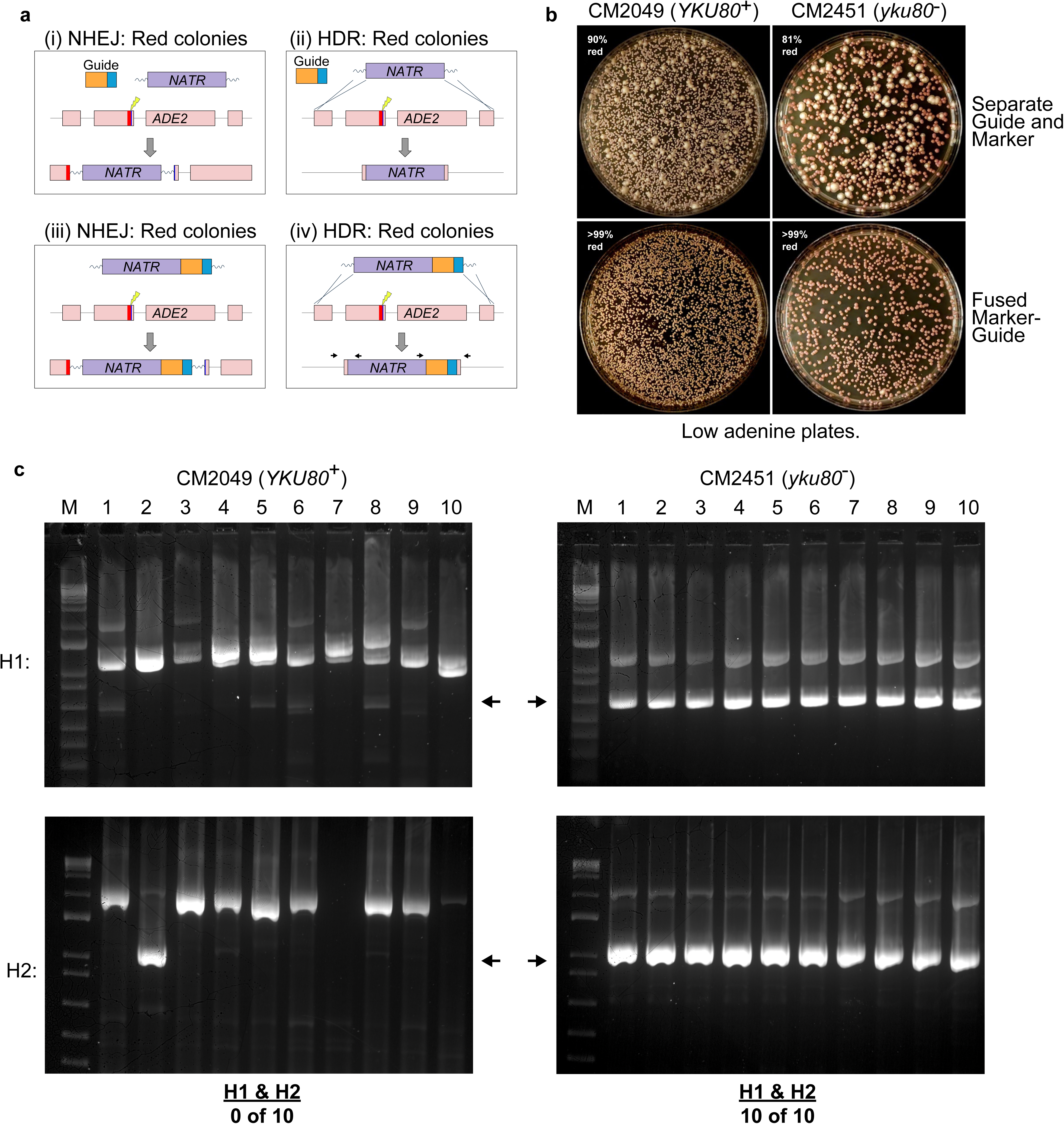
**Fused marker-guide in Ku-deficient background enhances HDR** (a) Diagrams of possible transformation outcomes. (i) A separate guide molecule targets Cas9 to cut *ADE2*, and NHEJ causes insertion of the separate marker molecule, disrupting *ADE2*, which yields red colonies. (ii) Similar to (i), but HDR leads to deletion of *ADE2* and red colonies. (iii) With a fused marker-guide, NHEJ causes marker-guide insertion and yields red colonies. (iv) As in (iii), but HDR deletes *ADE2* and yields red colonies. NOTE: Arrows in (iv) indicate primers used for junction PCR. (b) Top: When separate molecules of guide and marker were transformed into either *YKU80*^+^ or a Ku-deficient background, 81-90% of transformants were mutants of *ADE2*. Bottom: When a fused marker-guide was transformed into either background, greater than 99% of transformants were mutants of *ADE2*. (c) Junction PCRs of 10 red colonies from each background transformed with fused marker-guide indicate that HDR at both ends is rare (0 of 10) with *YKU80*^+^, whereas HDR at both ends is frequent (10/10) in a Ku-deficient strain. NOTE: Arrows indicate the expected band size for an HDR event.

In the case of the marker without homology arms, very few transformants were recovered (BOUCHER *et al*. 2024). In contrast, numerous colonies grew on the selective plate when a marker with homology arms was used (as in Fig. 1b, top left panel). Additionally, most colonies were red, indicating that a mutation of *ADE2* was produced (BOUCHER *et al*. 2024). These results suggest that without Ku80, Cas9-expressing *C. neoformans* is virtually unable to survive a double-strand DNA break programmed by a sgRNA-encoding DNA unless a template with homology to regions flanking the Cas9/sgRNA cut site is also introduced.

### Fused marker-guide system enables extremely efficient HDR in Ku-deficient strains

While the *ADE2* transformations indicated that *C. neoformans* might be strongly biased to HDR in the absence of functional Ku80, the overall efficiency of inducing a mutation in *ADE2* was still below 100% (Fig. 1b top panels). We suspected some transformants corresponding to white colonies received the marker without a guide. The lack of a guide could mean 1) *ADE2* was not cut, 2) an off-target integration or 3) maintenance of the marker extrachromosomally. Any of these possibilities might allow these cells to grow on selective media as white colonies (Ade^+^) rather than red colonies (Ade^-^) (Fig. 1b top panels). Therefore, we next performed transformations using linear DNA molecules that encoded both a selectable marker and a sgRNA flanked by targeting homology. Such molecules were assembled using fusion PCR (HUANG *et al*. 2022). We expected that transformation of such fused marker-guide DNAs into Cas9-expressing *C. neoformans* would limit the possibility of cells maintaining the marker extrachromosomally. Indeed, we observed that virtually all colonies were red after transformation and selection (Fig. 1b, bottom panels), regardless of the presence of Ku, indicating that the *ADE2* gene was mutated in each transformant. To determine the frequency of accurate HDR in Ku80^+^ vs. ku80^-^ backgrounds, we selected 10 red colonies from each transformation and performed junction PCR spanning each homology arm. In the Ku80^+^ background, 0 of 10 colonies had acquired both junctions, but in the ku80^-^ background, 100% (10 of 10) colonies displayed both junctions (Fig. 1c), demonstrating the substantial improvement of HDR frequency in this strain.

The experiments described above with *ADE2* showed that nearly 100% of transformants contained a mutation in the targeted gene when marker and guide were fused and flanked by homology arms, regardless of genetic background. However, these results are limited to one gene and a relatively small sample size. To more accurately determine the relative levels of HDR in these strains in another context, we set up a similar set of transformations to tag the *TKL1* gene (encoding transketolase; CNAG_07445) with mNeonGreen (mNG) (SHANER *et al*. 2013) by homologous recombination using a DNA construct harboring mNG, a transcriptional terminator, a sgRNA-encoding segment, and a nourseothrycin resistance (*NATR*) marker (Fig. 2a). Because *TKL1* expression is high, and mNG expression requires precise homologous recombination, flow cytometry can easily distinguish cells that had undergone HDR from those that had not. For these experiments, we selected transformants in a pool and then examined the fraction of drug-selected cells that expressed mNG using flow cytometry.

**Figure 2.**
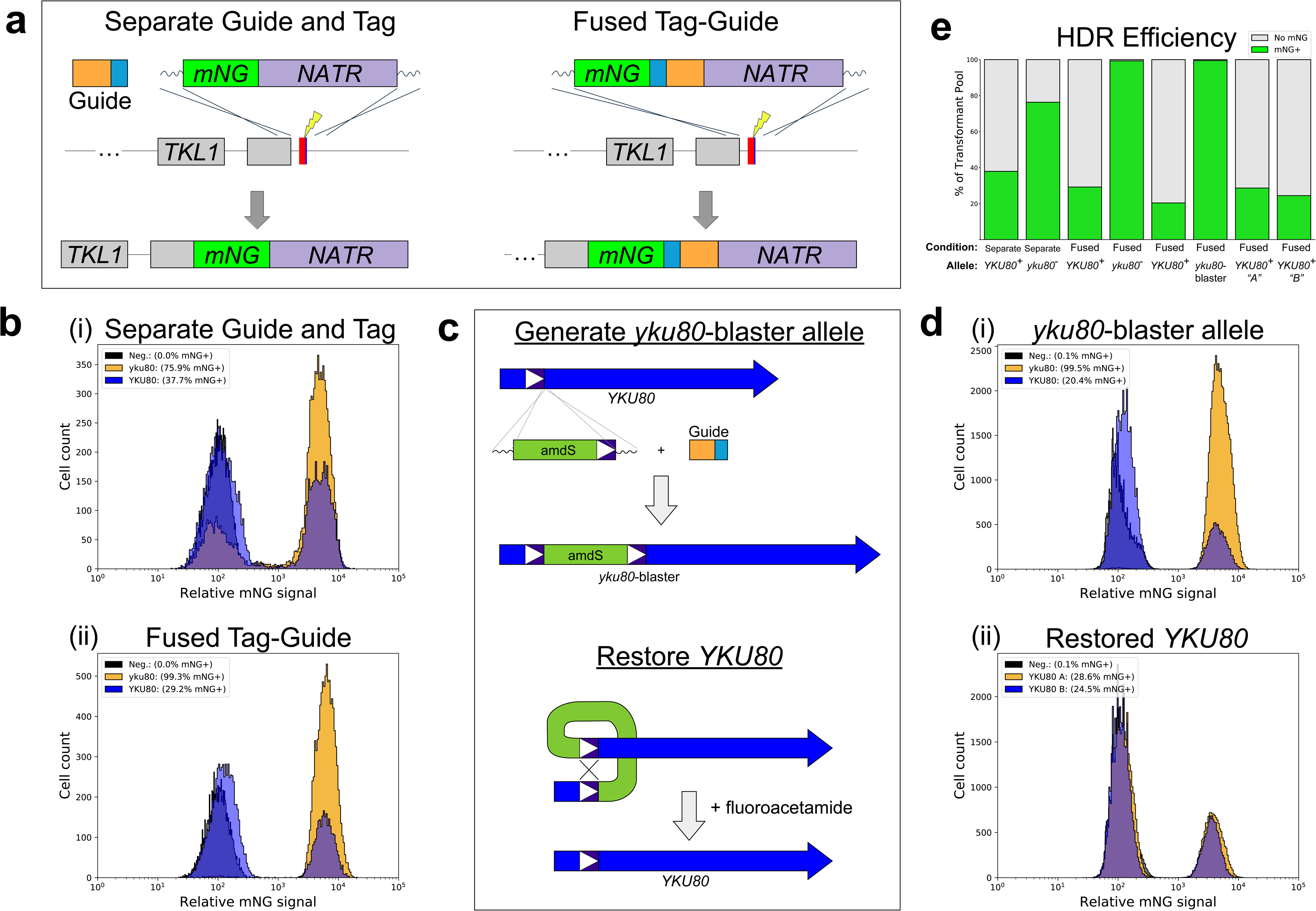
**Fused marker-guide in Ku-deficient backgrounds yields greater than 99% HDR** (a) Diagrams of the two approaches to tagging (separate guide and tag vs. fused tag-guide). In each case, a cut is made in the 3’UTR of *TKL1*, and after HDR the stop codon and target site are eliminated. (b) Flow cytometry results comparing *YKU80*^+^ and Ku-deficient (*yku80*::*NEOR*) backgrounds showing that the combination of fused tag-guide with Ku deficiency generates a transformant pool with over 99% mNG-positive cells, which are the result of HDR. (c) Diagrams showing (top) the transformation strategy to create the Ku-deficient strain *yku80*- blaster and (bottom) that *YKU80* can be restored by selecting for rare spontaneous recombinants that have lost the *amdS* marker. The 313 bp blaster sequence spans exons 1-3 (L23 to S92). NOTE: Introns are excluded from *YKU80* for simplicity. (d) Flow cytometry results comparing *YKU80*^+^ and *yku80*-blaster backgrounds showing that this Ku-deficient strain can also generate over 99% mNG-positive cells, and that after selection against *amdS*, two independent isolates return to HDR frequencies similar to *YKU80*^+^. (e) Summary of flow cytometry data in panels (b) and (d).

We compared separate guide and mNG-marker (separate guide and tag) vs. fused mNG-marker-guide (fused tag-guide). In each case, the mNG-containing donor DNA had homology arms (Fig. 2a). After transformation with two DNA molecules, a separate guide-encoding molecule and separate tag-encoding molecule, the Ku80^+^ background yielded only 37.7% mNG^+^ cells, while the ku80^-^ background yielded 75.9% mNG^+^ cells (Fig. 2b(i)). In the Ku80^+^ background, transformation with a fused tag-guide construct did not increase the fraction of the selected population that was mNG^+^—it was only 29.2%, presumably because NHEJ dominantly programs the outcomes for this particular gene/construct (Fig. 2b(ii)). However, in the transformation with fused tag-guide molecule harboring targeting homology in the ku80^-^ background, over 99% of nourseothrycin-resistant cells were mNG^+^, indicating that lack of Ku80 together with the fused tag-guide approach boosts HDR efficiency to over 99% (Fig. 2b(ii)).

### Generation and use of a reversible mutation in *YKU80*

Although the ku80^-^ strain allowed us to perform transformations using short homology arms with near-perfect HDR efficiency, the aforementioned phenotypes of a ku80^-^ mutant limit the utility of this approach. To minimize these drawbacks, we engineered a “*yku80*-blaster” allele, in which a mutation of *YKU80* is reversible. As before, this strain also expresses a codon-optimized Cas9 protein. In this strain, *YKU80* is interrupted with an insertion containing the acetamide utilization *amdS* marker (ERPF *et al*. 2019) flanked by a 313 bp direct repeat derived from the *YKU80* gene (Fig. 2c, top). The mutation in the genome is designed such that rare spontaneous recombination between the direct repeats produces a wild-type *YKU80* allele (Fig. 2c, bottom). Because *amdS* expression produces sensitivity to the toxic acetamide analog fluoroacetamide (FA) (ERPF *et al*. 2019), growth on FA should, in principle, select for the restoration of *YKU80* via recombination between the direct repeats.

To test these ideas, we again used flow cytometry and tagging of the *TKL1* gene with mNG as a testbed. First, we performed transformations to compare the wild-type *YKU80* allele vs. the *yku80*-blaster allele using fused tag-guide. While the Ku80^+^ background yielded a minority of mNG^+^ cells (20.4%), the transformant pool from the *yku80-*blaster background displayed almost all mNG^+^ cells (99.5%) [Fig. 2d(i)]. Next, we tested whether Ku function could be restored in the parental *yku80*-blaster strain. We selected against *amdS* using FA and chose two FA-resistant isolates for analysis. When we transformed these *YKU80*-restored isolates (A and B) with the *TKL1-mNG* fused marker-guide construct, we observed the percentage of mNG^+^ transformed cells to be 28.6% and 24.5%, respectively, similar to what we observed in the Ku80^+^ background (Fig. 2d(ii)). These results are summarized in Fig. 2e. To confirm the restoration of Ku80, we performed PCR with primers flanking the site of insertion of the *amdS* gene. This yielded a band consistent with the desired recombination between the direct repeats. Sanger sequencing of the PCR product obtained the expected *YKU80* sequence (including two engineered silent SNPs). We conclude that the *yku80*-blaster strain can be reverted to wild-type *YKU80*.

To test the effectiveness of the *yku80-*blaster strain on another gene, we also targeted *URA5* with either a fused marker-guide DNA molecule flanked by short homology arms or a fused marker-guide DNA molecule lacking homology arms. Transformation of the *yku80-*blaster strain was compared to the precursor *YKU80*^+^ strain as well as the two *YKU80*-restored isolates (A and B) [Fig. 3a]. Transformants were selected on nourseothrycin plates. As with *ADE2* in the *yku80Δ* (BOUCHER *et al*. 2024), transformants were obtained in the *yku80-*blaster background only when homology arms were present in the transformed construct. Importantly, the *YKU80*-restored isolates also yielded transformants without homology arms in the marker- guide donor DNA (Fig. 3b), further confirming restoration of Ku function in these backgrounds.

**Figure 3.**
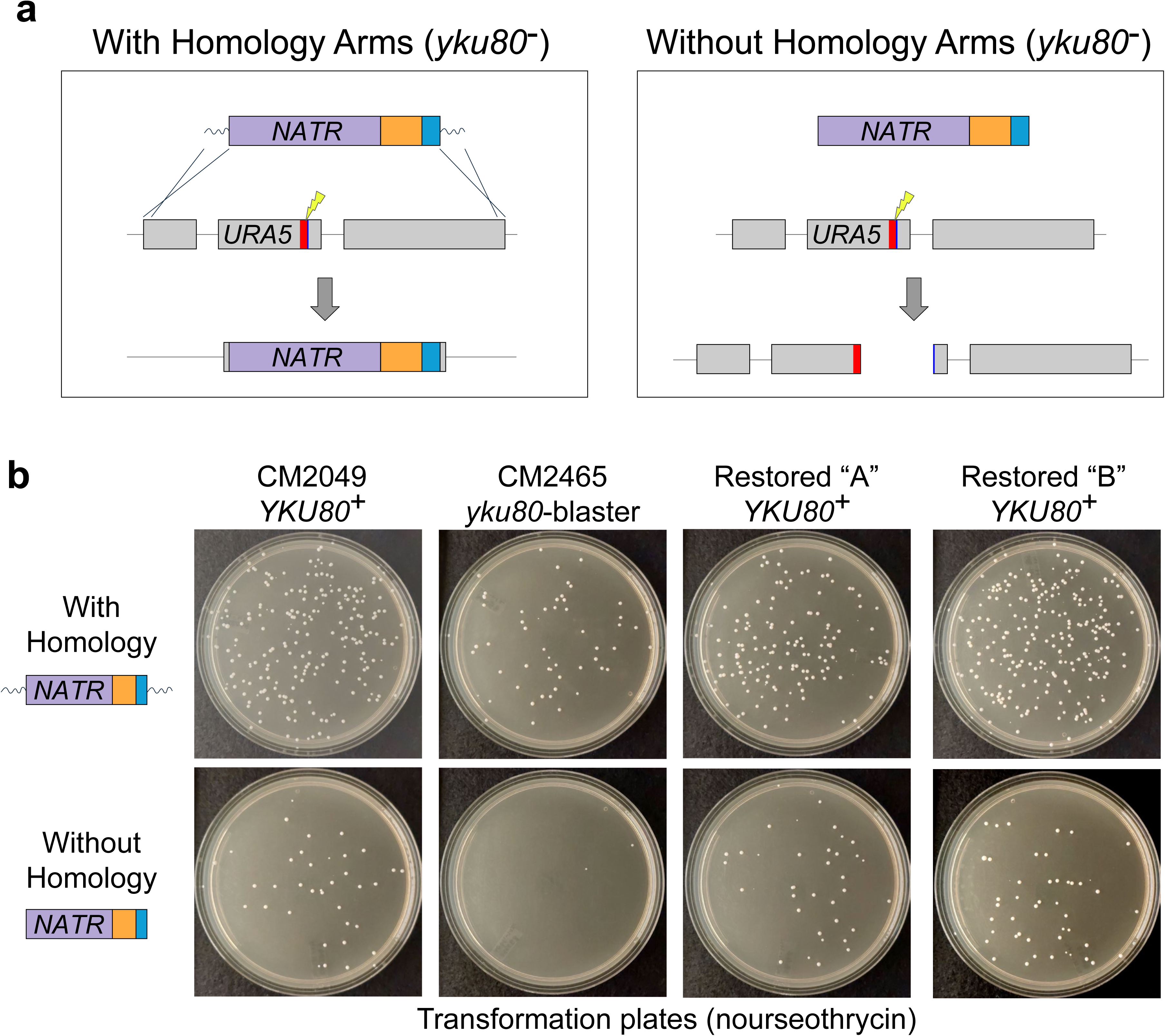
**Mutating *URA5* with the *yku80*-blaster strain** (a) Diagrams showing the possible outcomes of transformations in a Ku-deficient strain. Left: A fused marker-guide with homology arms yields *URA5* deletion via HDR. Right: Without homology arms, the marker-guide breaks *URA5* but cannot repair the cut, which leads to cell death. (b) When targeting *URA5* using a fused marker-guide flanked with homology arms, transformants are obtained in all tested backgrounds (*YKU80*^+^, *yku80*-blaster, and both restored *YKU80*^+^ isolates). Without homology arms, a fused marker-guide fails to yield transformants in the *yku80*-blaster strain, confirming its NHEJ deficiency; similar numbers of transformants among the *YKU80*^+^ background and both restored *YKU80*^+^ isolates indicate a functional NHEJ repair pathway.

### Plasmid series for constructing marker-guide DNAs for genome manipulation

Finally, we constructed a series of plasmids that enable facile generation of sgRNA- marker constructs for both gene deletion and C-terminal tagging in *C. neoformans*. These have been deposited to Addgene. The first set of plasmids contains marker-guide fusions, including the markers *HYGR*, *NATR*, *NEOR*, and *amdS*. In each plasmid, the marker is followed by U6 promoter, a stuffer sequence with BplI restriction site, and the sgRNA scaffold (Fig. 4a). Similarly, we constructed fused mNG-guide-marker plasmids for C-terminal tagging (Fig. 4b). A fused marker-guide or tag-guide-marker DNA can be generated by one of two methods. A 20 bp target sequence can replace the stuffer sequence via fusion PCR, as described in the Methods. Alternatively, cloning can be performed as follows: after digesting the plasmid with *Bpl* I, the break can be bridged with a 60 bp ssDNA oligo containing the desired 20 bp target sequence flanked by U6 and sgRNA scaffold homologies (see NEB #E2621). Note that because the *amdS* gene also contains a *Bpl* I recognition site, its use is limited to the fusion PCR method.

**Figure 4.**
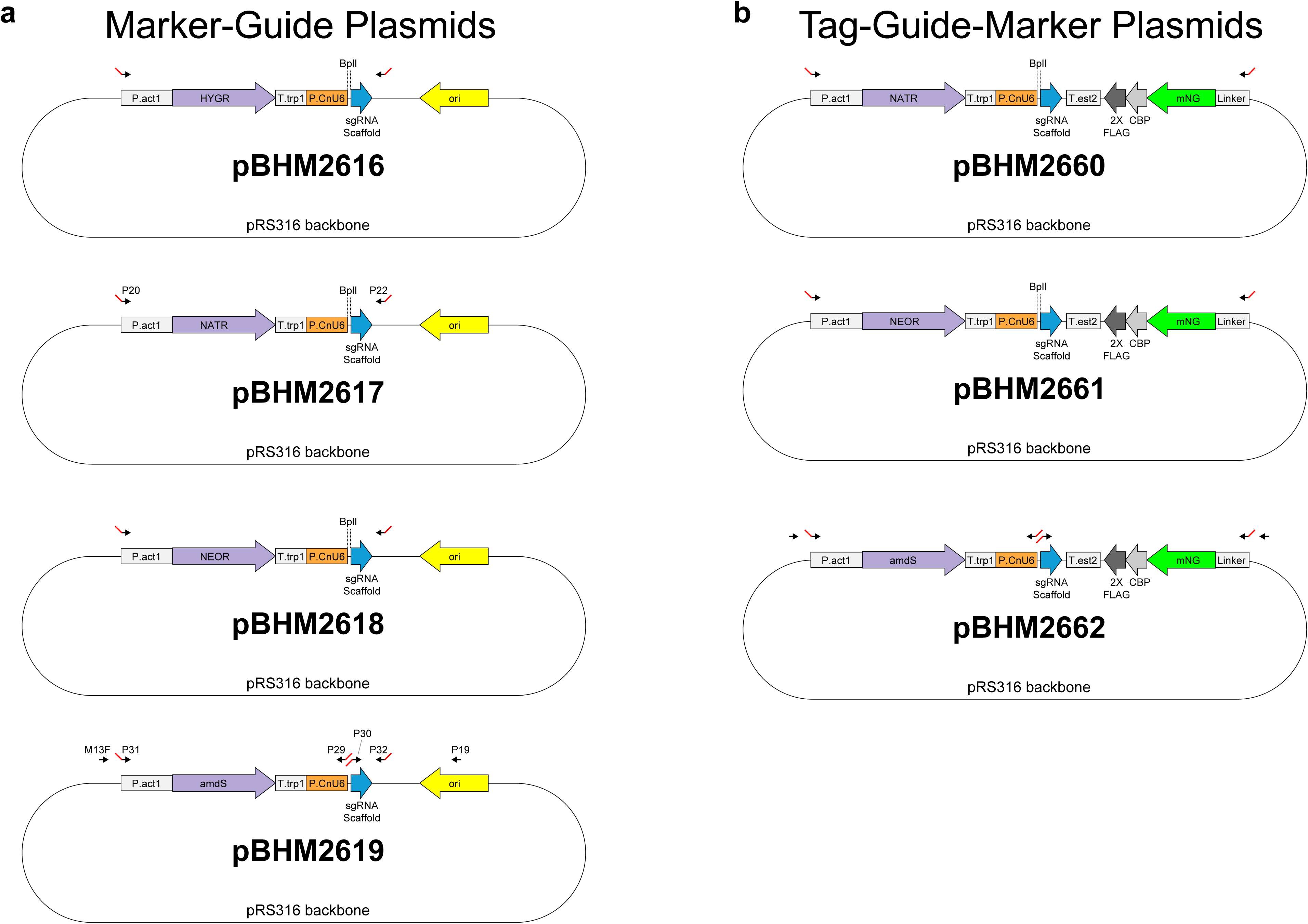
**Plasmid maps** (a) Marker-guide plasmids for *HYGR*, *NATR*, *NEOR*, and *amdS* (pBHM2616-2619). Each marker is driven by the *ACT1* promoter and terminated by the *TRP1* terminator. The sgRNA expression is driven by the U6 promoter and terminated by 6 Ts (not shown); a stuffer sequence includes the *Bpl* I restriction site for cloning. Arrows indicate primers used for fusion PCR with red tails indicating short homologies where applicable. As examples: pBHM2617 specifies P20 and P22, which were used for the *ADE2* deletion cassette with homology arms; pBHM2619 includes primers for fusion PCR to generate a *URA5* marker-guide deletion cassette with homology arms (Step 1A: M13F + P29, Step 1B: P30 + P19, and Step 3: P31 + P32). (b) Tag-guide-marker plasmids for *NATR*, *NEOR*, and *amdS* (pBHM2660-2662). Each marker- guide portion is as described in (a), and the tag portion includes a linker followed by codon- optimized mNeonGreen (mNG) as well as CBP and 2xFLAG (separated by short linkers; not shown) with a terminator from *EST2*. Note that because the *amdS* gene also contains a *Bpl* I recognition site (not shown), its use is limited to the fusion PCR method.

## Discussion

Exploiting homology-directed repair (HDR) in *C. neoformans* to edit the genome of this important human fungal pathogen has been a powerful and widely used technique in the field for over thirty years (TOFFALETTI *et al*. 1993). While the shift from laborious biolistic transformation with long homology arms to the use of electroporation of Cas9-expressing strains with short homology arms (HUANG *et al*. 2022) has eased the burden of generating targeting constructs and mutants, the efficiency is variable because *C. neoformans* maintains an NHEJ repair bias. Ku-defective mutants can enhance HDR efficiency in *C. neoformans* (GOINS *et al*. 2006; LIN *et al*. 2015; HUANG *et al*. 2022), although its efficacy has been questioned (FAN AND LIN 2018). Furthermore, *C. neoformans* Ku-defective mutants are known to have increased mutation rates (JUNG *et al*. 2021) as well as reduced fitness in the host and under stress conditions (LIU *et al*. 2008; JUNG *et al*. 2021), limiting the attractiveness of their use.

We show here that such limitations can be overcome. To tip the scale entirely towards HDR over NHEJ, electroporation of a fused marker-guide DNA flanked with short homology arms into a Ku-defective, Cas9-expressing background can yield over 99% HDR events. We show that extremely high HDR efficiencies are achieved when targeting three different genes: *ADE2*, *TKL1*, and *URA5*. Although this high efficiency is desirable, the drawbacks of a Ku- defective background limit its utility. Therefore, we created a Cas9-expressing *yku80::amdS* ”blaster” strain, which also achieves very high HDR efficiency, while enabling reversion to a Ku80^+^ phenotype by counter-selection using the *amdS* marker and fluoroacetamide. Not only does this ability to revert back to Ku80^+^ alleviate the issues associated with Ku-defective mutants, it also enables re-use of the *amdS* marker. Taken together, the methods and strains described here significantly accelerate *C. neoformans* molecular genetics.

## Methods

### Strains and Media

Strains were stored at -70°C in 15-25% glycerol. YPAD media (2% Bacto Peptone, 2% dextrose, 1% yeast extract, adenine at 120 mg/mL) with 2% agar was used to recover yeast stocks on plates or grow cultures in liquid at 30°C on a roller drum at 60 RPM (up to 10 mL) or in a shaker at 200 RPM (more than 20 mL). LB media with 2% agar was used to recover bacteria stocks on plates or grow cultures in liquid at 37°C on a roller drum at 60 RPM. For transformants harboring drug resistance (*NATR* or *NEOR*) markers, YPAD plates supplemented with clonNAT at 75 ug/mL or G418 at 100 ug/mL, respectively, were used for selection. Transformants with the *amdS* cassette were selected on acetamide plates (YNB without amino acids or ammonium sulfate, 2% glucose, 5 mM acetamide), and spontaneous loss of the *amdS* cassette was selected on fluoroacetamide plates (YNB, 10 mM ammonium sulfate, 2% glucose, 10 mM fluoroacetamide) (ERPF *et al*. 2019).

### Plasmid Construction

Plasmids pBHM2616 (*HYGR-empty sgRNA*) and pBHM2617 (*NATR-empty sgRNA*) were generated as follows. Three segments were amplified for each vector with overlapping homology between sequential products. All PCRs were performed using Q5 Polymerase (NEB) per the manufacturer’s instructions. The markers were amplified with primers P01 and P02 using either pSDMA25 or pSDMA57 as templates. The U6 promoter was amplified using primers P03 and P04, and the scaffold was amplified using P05 and M13R with pBHM2329 as template. Overlapping homology between these two segments also carried a sequence to introduce a *Bpl* I site to facilitate downstream sgRNA cloning. Fragments were assembled using gap repair in *S. cerevisiae* BY4741 into a pRS316 backbone using the lithium-acetate transformation protocol as described elsewhere (GIETZ AND SCHIESTL 2007).

Plasmids pBHM2618 (*NEOR-empty sgRNA*) and pBHM2619 (*amdS-empty sgRNA*) were constructed similarly. Marker cassettes for *NEOR* and *amdS* were each amplified with homology to pBHM2617’s P.act1 region and its CnU6 promoter using primers P06 and P02 with either pSDMA57 or pPEE7, respectively, as the template. Plasmid pBHM2617 was digested with *Not* I and *Sfo* I to remove the *NATR* cassette. Purified linear backbone and either the *NEOR* cassette or the *amdS* cassette were assembled in *S. cerevisiae* BY4741 as described above.

Plasmids pBHM2660-2662 (mNG-guide-markers) were constructed by inserting mNG flanked by a linker and CBP-2xFLAG into each marker-guide plasmid (pBHM2617-2619). Briefly, the marker-guide plasmid was digested with *Hind* III to cut downstream of the sgRNA scaffold and its 6T terminator. The linker-mNG-CBP-2xFLAG cassette was amplified from pBHM2404 with primers P07 and P08, which contain homology to the cut marker-guide plasmid. The purified linear backbone and the mNG cassette were assembled in *S. cerevisiae* BY4741, as described above.

Plasmid pBHM2659 (*amdS-yku80-blaster*): Two fragments were amplified with overlapping homology. From *YKU80* (CNAG_03637), 313bp was amplified using primers P09 and P10 with KN99ɑ gDNA as the template. The *amdS* cassette was amplified with primers P11 and P12 using pPEE7 as the template. Plasmid pRS316 was digested with BamHI and EcoRI to linearize the backbone. The fragments were assembled in *S. cerevisiae* BY4741 as described above.

After assembly of each plasmid in BY4741, yeast plasmid extract was prepared from 10-20 colonies using the Zymo Yeast Plasmid Miniprep II kit, and 1 ul was transformed into electrocompetent DH5ɑ *E. coli*. Plasmids were isolated from single colonies by miniprep.

All plasmid sequences were verified by long-read sequencing (Plasmidsaurus, Inc.).

### Electroporation

All electroporations were performed as described previously (HUANG *et al*. 2022). Except for transformations to tag *TKL1*, electroporated cells were spread on selective plates. For *TKL1* tagging, after recovery for 2 hours in 1 mL YPAD, transformation pools were grown for 40 hours in 20 mL YPAD with clonNAT at 75 ug/mL.

### Strain Construction

Strain CM2451 (*yku80*::*NEOR*). To disrupt *YKU80*, KN99ɑ with integrated Cas9 (CM2049) was electroporated with 500 ng of a guide targeting *YKU80* and 1 ug of a *NEOR* cassette. Transformants were selected on YPAD+G418 plates.

To delete (or disrupt) *ADE2*, electroporations were performed with conditions as follows. CM2049 or CM2451: 700 ng of guide targeting *ADE2* + 1 ug of a *NATR* cassette with ∼50bp homology arms. In CM2049 or CM2451: 1 ug of fused *NATR*-guide with ∼50bp homology arms. After recovery in YPAD, transformants were selected on nourseothrycin plates.

To test Ku80^+^ vs. ku80^-^ by tagging *TKL1* with mNG, electroporations were performed with conditions as follows. In CM2049 or CM2451: 700 ng of guide targeting *TKL1* + 1 ug of a mNG-*NATR* cassette with ∼50bp homology arms. In CM2049 or CM2451: 1 ug of fused mNG- *NATR*-guide with ∼50bp homology arms.

Strain CM2465 (*yku80*-blaster). To disrupt *YKU80*, KN99ɑ with integrated Cas9 (CM2049) was electroporated with 100 ng of a guide targeting *YKU80* and 750 ng of an *amdS*- blaster cassette, which included homology arms to the regions of *YKU80* immediately upstream and downstream of the cut site (P13 and P14). Transformants were screened via PCR for the correct junctions, indicating both 5’ and 3’ HDR. Sanger sequencing of each junction verified the presence of the expected sequences, including two engineered silent SNPs (S91 TCC-to-TCA and S92 TCA-to-TCC) to break the guide target site.

To test Ku80^+^ vs. the *yku80*-blaster allele by tagging TKL1 with mNG, electroporations were performed with conditions as follows. In CM2049 or CM2465: 1 ug of fused tag-guide with ∼50bp homology arms. After recovery and selection, transformant pools were diluted to O.D. ∼0.1 in 5 ml YNB and grown for 3 – 4 hours. For flow cytometry, 200 ml of the log phase culture is added to 2 – 3 ml of YNB and transferred to a 5 ml flow cytometry tube.

Strains CM2494 and CM2495 (YKU80 Restored “A” and “B”): To restore Ku80 function, 5 mL YPAD cultures of CM2465 were grown overnight to OD ∼1.0. Approximately 1 million cells were plated on fluoroacetamide (10 mM) plates. Three days later, colonies were patched to fresh fluoroacetamide plates. The following day, we performed patch PCR with primers P15 and P16 flanking the 313bp blaster region of *YKU80*. Patches that yielded the expected 560bp amplicon were re-streaked to single colonies, and two independent isolates were selected to save glycerol stocks and prepare genomic DNA. The PCR spanning the *YKU80* blaster region was repeated, and the 560bp amplicon was Sanger sequenced to confirm accurate HDR of *YKU80*.

To test the restored Ku80 function (after selection against *amdS* in *yku80*-blaster strain background) by tagging *TKL1* with mNG, electroporations were performed as follows. In CM2049, CM2494, or CM2495: 2 ug of fused tag-guide with ∼50bp homology arms. After recovery and selection, transformant pools were diluted to O.D. ∼ 0.1 in 5 ml YNB and grown for 3 – 4 hours. For flow cytometry, 200 ml of the log phase culture was added to 2 – 3 ml of YNB and transferred to a 5 ml flow cytometry tube.

### Fusion PCR

PCR steps to generate guides are as follows. Using pBHM2329 as a template, part A was amplified using the M13F primer paired with a reverse primer, which included a 20bp overhang corresponding to the reverse complement of the desired guide sequence (e.g. for *ADE2*, P18). Part B was amplified using a forward primer that includes a 20bp overhang corresponding to the desired guide sequence (e.g. for *ADE2*, P17) and the M13R primer. Step 1: Each part was amplified following Ex Taq’s recommended protocol with 1 ng plasmid template per 50 ul reaction and the following thermocycler conditions: 95°C for 3 minutes followed by 30 cycles (95°C for 30 seconds, 52°C for 30 seconds, and 72°C for 45 seconds) followed by 72°C for 5 minutes and held at 12°C. Part A and part B were combined and purified following the manufacturer’s protocol for PCR column cleanup using the NucleoSpin Gel and PCR Clean-up from Machery Nagel. Combined, cleaned A+B were then fused in Step 2. This reaction includes Ex Taq reagents as in Step 1 but in place of template and primers only purified parts A+B (10 ul per 50 ul reaction) were included. Step 2 thermocycler conditions: 95°C for 3 minutes followed by 10 cycles (95°C for 30 seconds, 55°C for 3 minutes, and 72°C for 3 minutes) followed by 72°C for 3 minutes and held at 12°C. Finally, the fused product was then amplified using primers M13F + M13R in Step 3. Thermocycler conditions: 95°C for 3 minutes followed by 30 cycles (95°C for 30 seconds, 52°C for 30 seconds, and 72°C for 45 seconds) followed by 72°C for 5 minutes and held at 12°C. PCR steps to generate a fused marker-guide are as described above with minor modifications. Each reaction included 5% DMSO, the template was pBHM2617 (*NATR*-empty sgRNA) or pBHM2618 (*NEOR*-empty sgRNA), the primer P19 was used in place of the M13R primer, extension times were increased to 1 minute/kb in Steps 1 and 3, and in Step 2 both annealing and extension times were increased to 5 minutes. When homology arms were needed for deletion cassettes, the M13F and M13R primers were exchanged for primers with the same annealing sequence at the 3’ end and appropriate homology extensions at the 5’ end. For example, *ADE2* deletion cassettes used primers P20 + P21 (*NATR*) or P20 + P22 (fused *NATR*-guide).

### Colony PCR

To prepare input for PCR, 1-5 ul of cells were pipetted to mix into 100 ul water in strip tubes. The cell mixture was flash frozen in liquid nitrogen for 5-10 seconds and immediately transferred to a preheated thermocycler block set to 100°C and boiled for five minutes. While cells boiled, a 2X master mix was prepared containing 400 mM Tris-HCl (pH 8.8), 40 mM MgSO_4_, 200 mM KCl, 200 mM (NH_4_)_2_SO_4_, 2% (v/v) Triton X-100, 10% DMSO, 400 uM dNTPs, 500 uM forward primer, 500 uM reverse primer, in house Taq polymerase at 10X, in house Pfu polymerase at 2X, and water. Boiled cells were allowed to cool at room temperature for one minute, and equal volumes of boiled cells and 2X master mix were pipetted to mix thoroughly. Leftover boiled cells were stored for subsequent PCRs at -20°C. Thermocycler conditions: 92.5°C for 3 minutes followed by 37 cycles of (92.5°C for 15 seconds, 45°C for 15 seconds, 72°C for 1 minute/kb) with a final extension at 72°C for 5 minutes.

### Genomic DNA Purification and Sanger Sequencing

Genomic DNA was prepared by bead beating and column purification. Briefly, 0.5 mL of a saturated YPAD culture was pelleted, washed once with 0.5 mL water, and pelleted again. The pellet was flash frozen in liquid nitrogen for 5-10 seconds and resuspended in 0.5 mL lysis buffer: 100 mM Tris, pH 8, 50 mM EDTA, and 1% SDS. This mixture was transferred to a screw-cap cryo-tube containing about 0.1 mL zirconia beads (0.5 mm) for bead beating on a table top vortexer at top speed for five minutes. Cell debris was pelleted by centrifugation at 20,000g for five minutes at 4°C, and 0.2 mL supernatant was transferred to a fresh 1.5 mL tube containing 0.4 mL DNA binding buffer from the Zymo 25 ug Clean & Concentrate kit. The kit protocol was followed with two washes, and DNA was eluted with 0.1 mL water. From this purified DNA, 2 ul was used for input per 50 ul PCR reaction following the manufacturers’ protocols for either Q5 or Ex Taq (including 5% DMSO). Amplicons were verified on agarose gels, column purified, and submitted for Sanger sequencing at Genewiz.

### Flow Cytometry

All cultures were grown in YNB to log phase (OD 0.3 - 0.4) unless otherwise stated. Samples were run on the BD FACSCelesta Multicolor Flow Cytometer until at least 10,000 events were recorded. Data was processed with the FlowCal python package to gate on live cells using the density2d function with parameters: FSC-A vs. SSC-A and gate_fraction = 0.95 followed by gating on single cells using the density2d function with parameters: FSC-A vs. FSC- H and gate_fraction = 0.95. Gated singlets were then analyzed with a custom python script, which normalized FITC-H (mNG) values against the negative control’s FITC-H median and generated plots with FlowCal v1.3.0, matplotlib v2.2.2, and seaborn 0.11.2 packages.

---

**Table 1:**
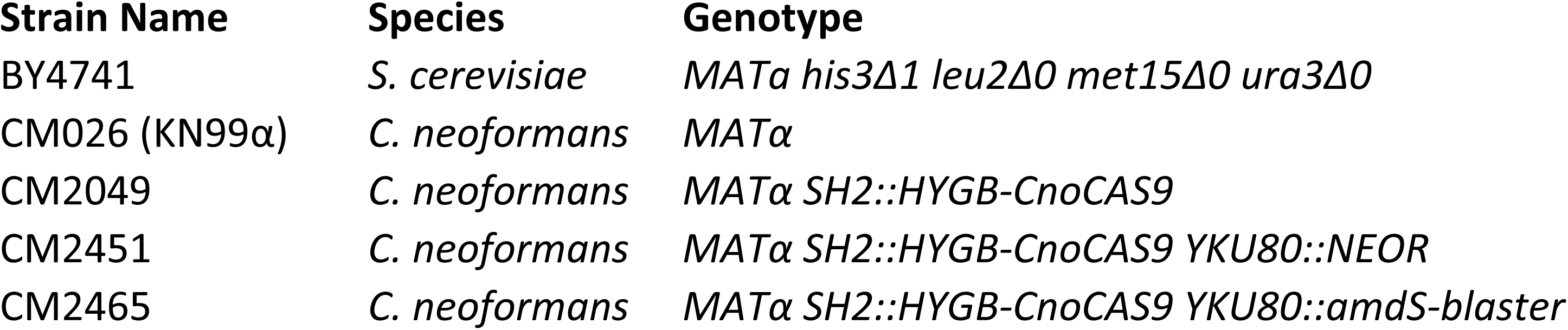
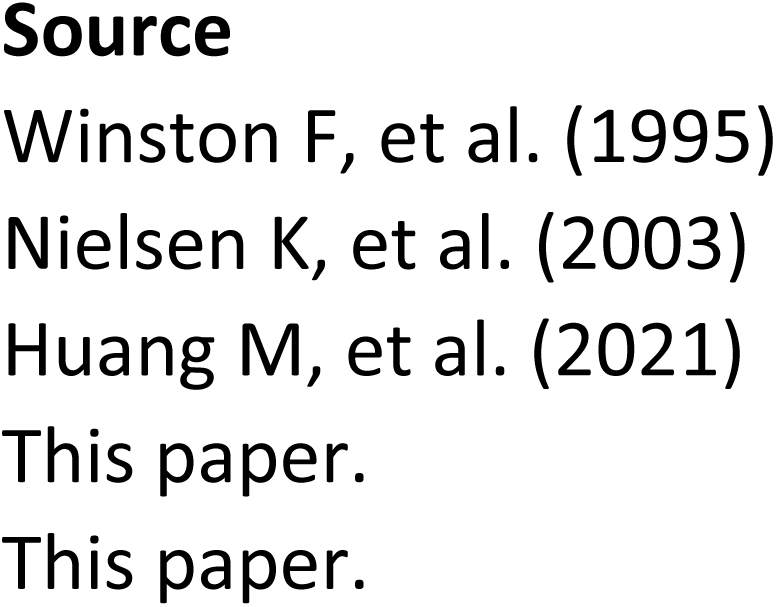
Strains.

**Table 2:**
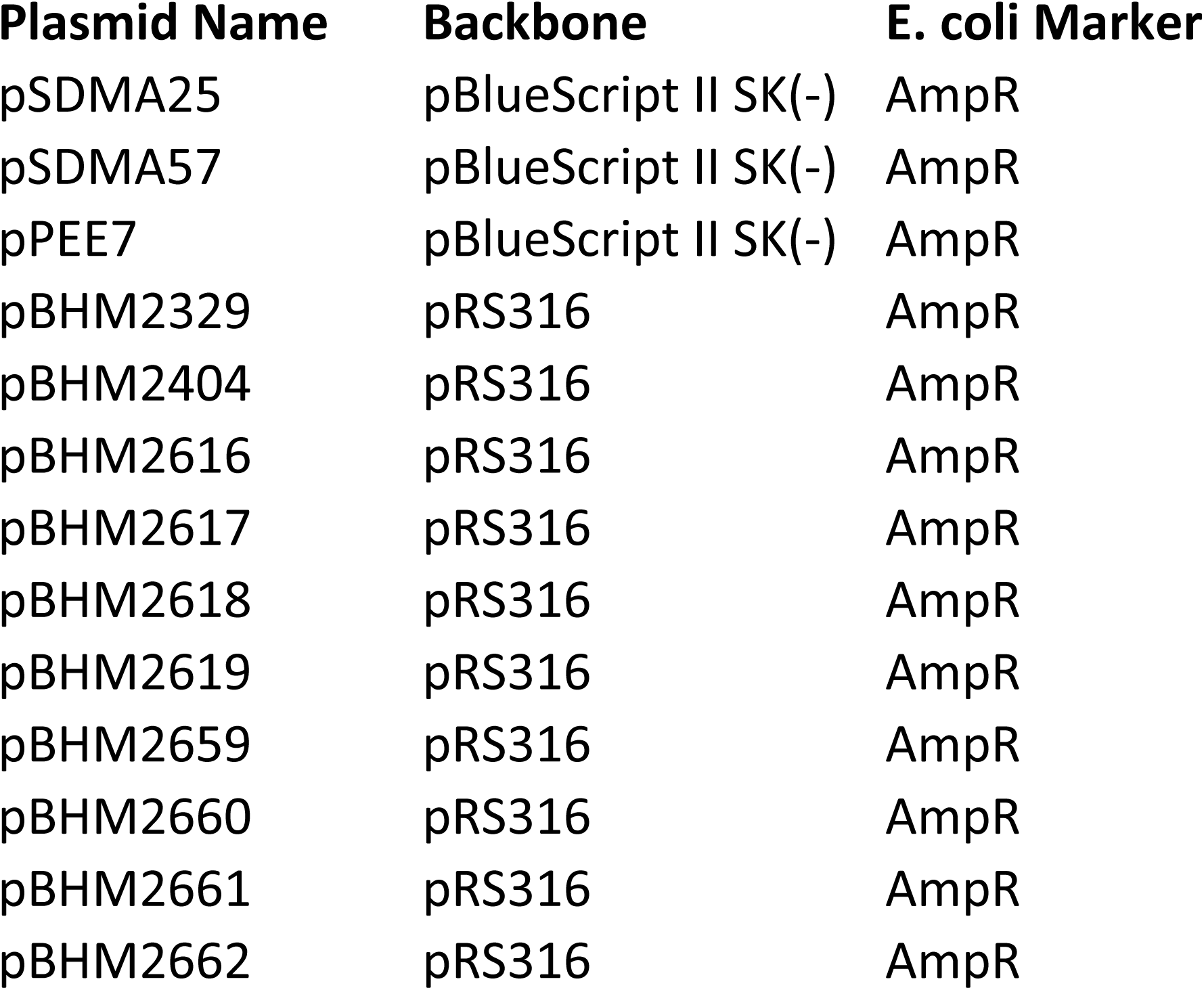
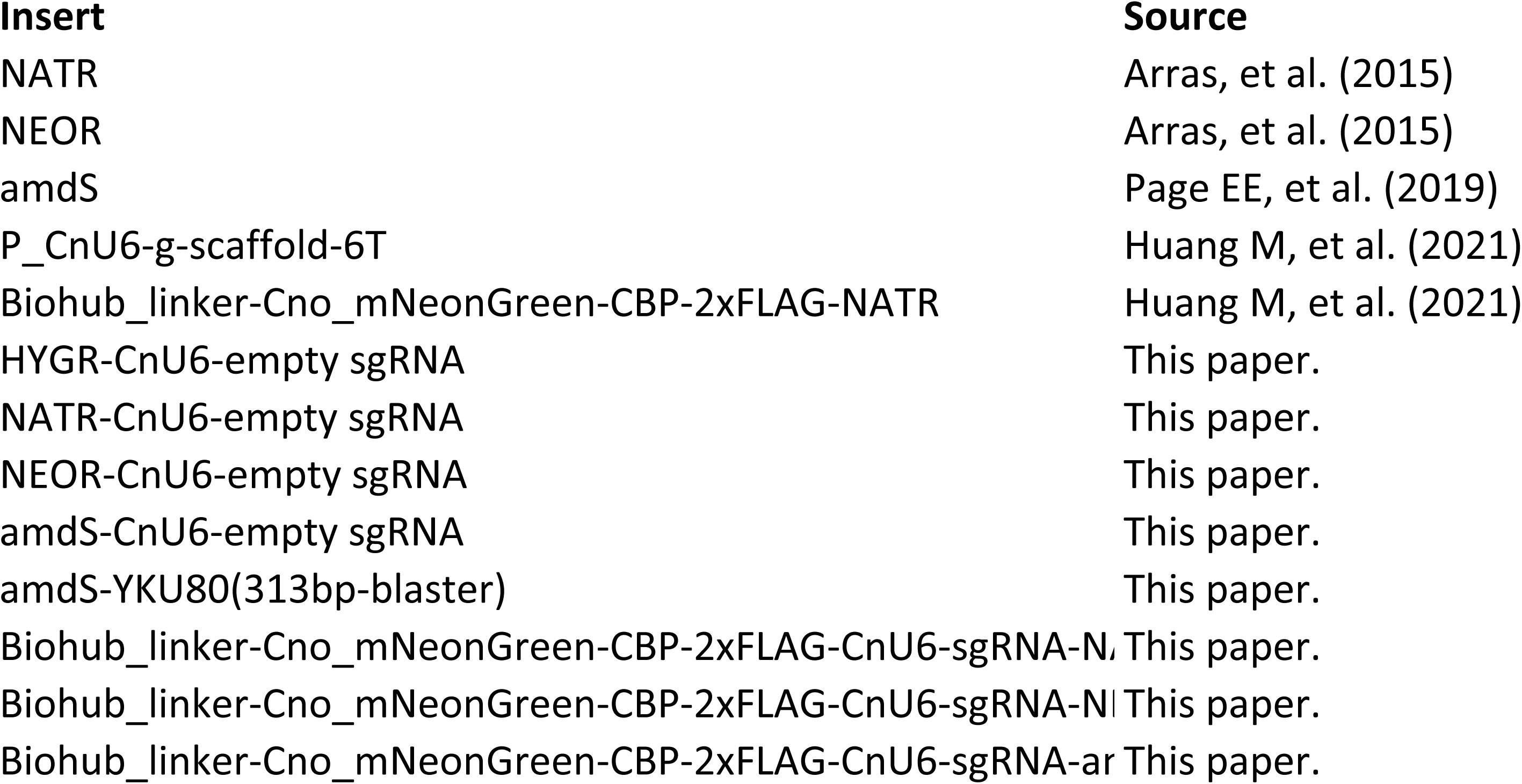
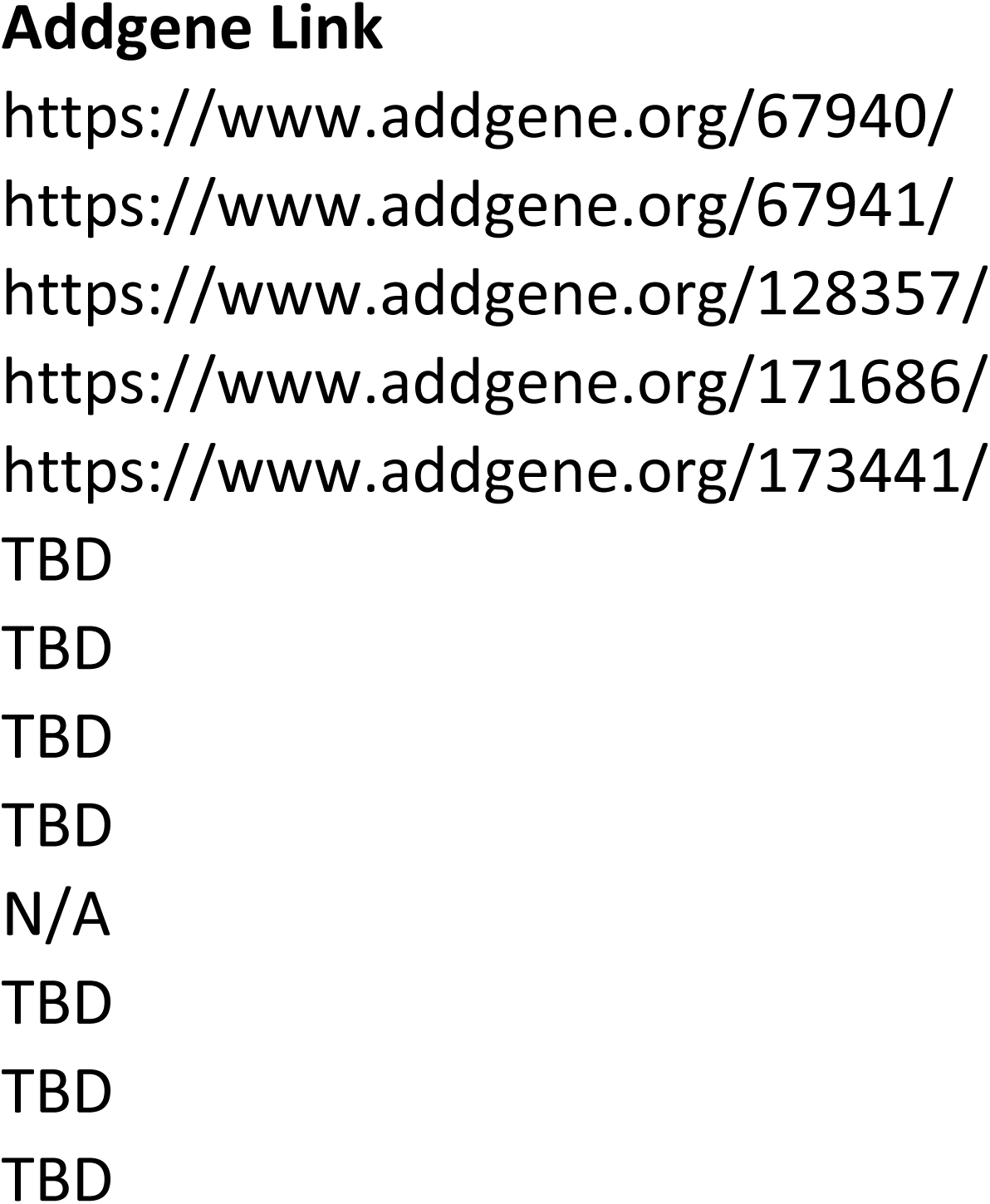
Plasmids.

**Table 3:**
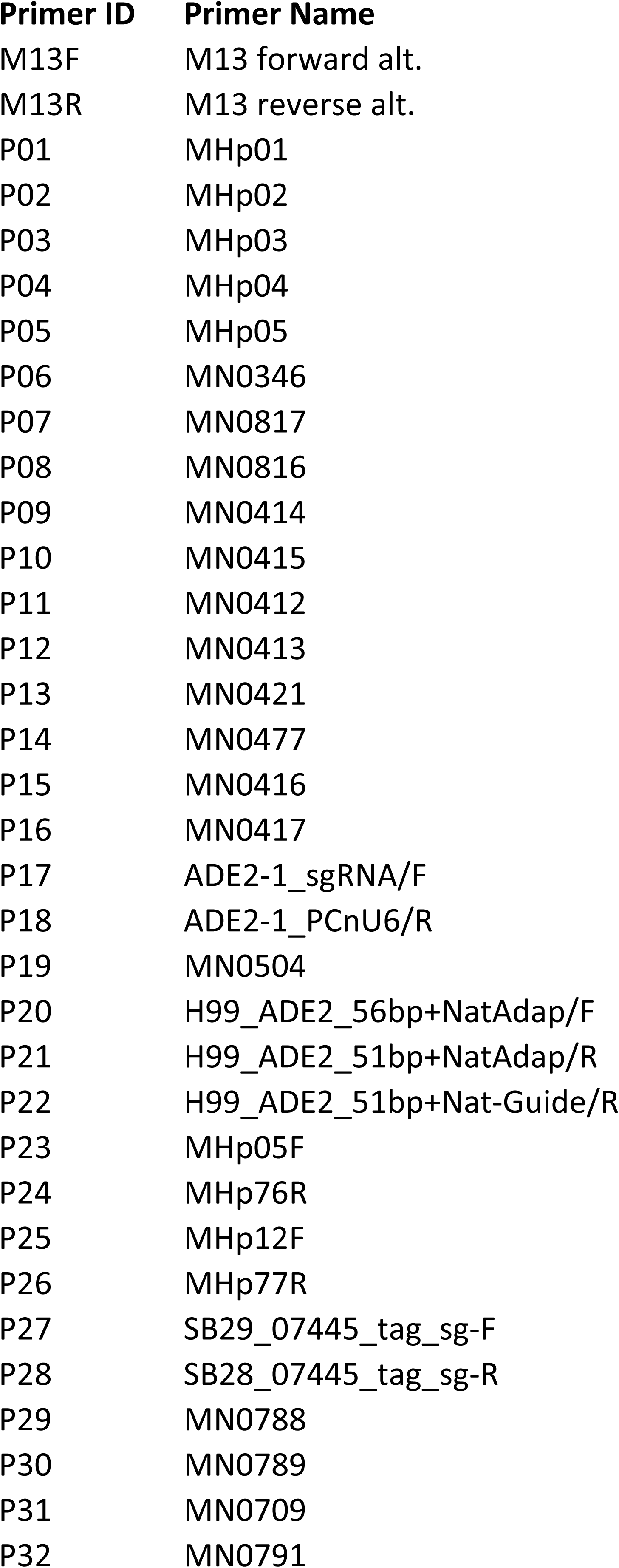
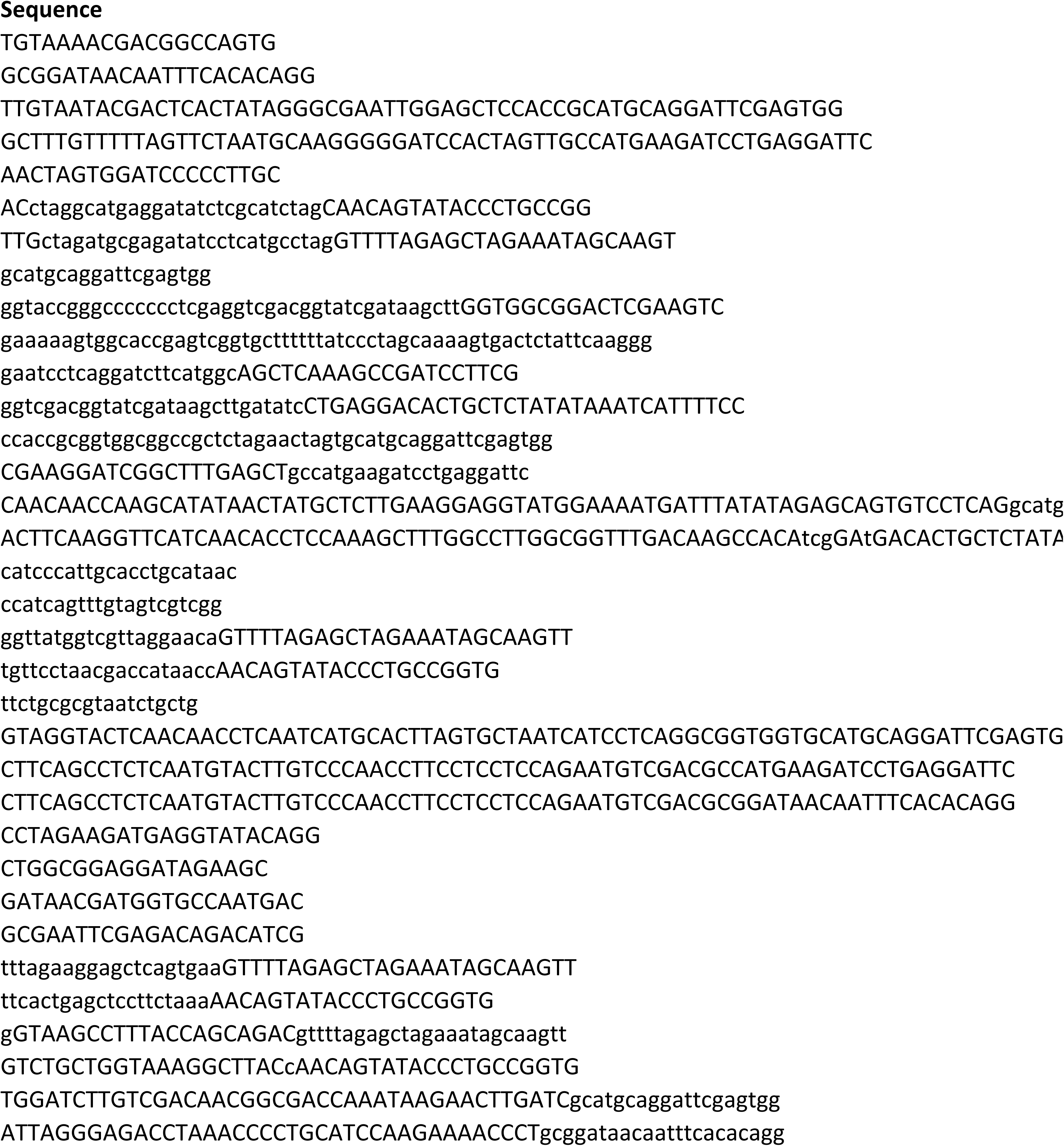
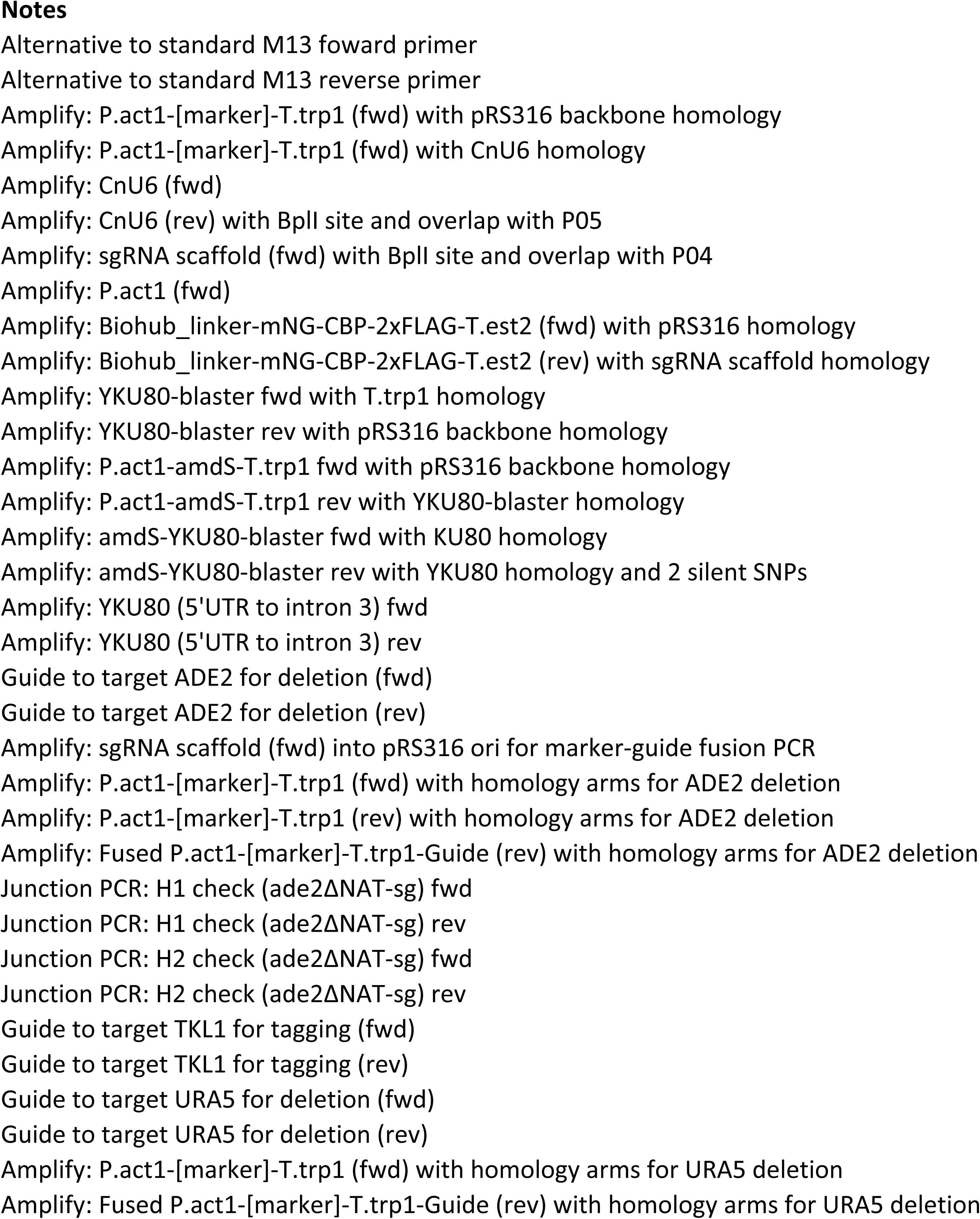
Primers.

